# Penetration of Topically Applied Polymeric Nanoparticles across the Epidermis of Thick skin

**DOI:** 10.1101/2023.08.16.553364

**Authors:** Andrea Antony, Gayathri Raju, Ahina Job, Meet Joshi, Sahadev Shankarappa

**Author notes:** Corresponding author /.

## Abstract

The natural barrier function of the epidermal skin layer poses a significant challenge to nanoparticle-mediated topical delivery. A key factor in this barrier function is the thickness of the stratum corneum (SC) layer within the epidermis, which varies across different anatomical sites. The epidermis from the palms and soles, for instance, have thicker SC compared to those from other areas. Previous studies have attempted to bypass the SC layer for nanoparticle penetration by using physical disruption; however, these studies have mostly focused on non-thick skin. In this study, we investigate the role of mechano-physical strategies on SC of thick skin for transdermal nanoparticle penetration. We characterize and compare two mechano-physical strategies, namely tape-stripping and microneedle abrasion, for epidermal disruption in both thick and thin skin. Furthermore, we examine the impact of SC disruption in thick and thin skin on the penetration of topically applied 100 nm sized polystyrene nanoparticles using an ex-vivo model. Our findings show that tape-stripping reduced the overall thickness of SC in thick skin by 87%, from 67.4 ± 17.3 µm to 8.2 ± 8.5 µm, whereas it reduced thin skin SC by only 38%, from 9.9 ± 0.6 µm to 6.2 ± 3.2 µm. Compared to non-thick skin, SC disruption in thick skin resulted in higher nanoparticle diffusion. Tape-stripping effectively reduces SC thickness of thick skin and can be potentially utilized for enhanced penetration of topically applied nanoparticles in skin conditions that affect thick skin.

## 1. Introduction

Topical mode of drug delivery remains a simple and highly patient compliant route that offers several benefits including convenience and pain-free administration of drugs. Especially with the advent of nanoparticle-based systems, there have been several improvements in terms of drug absorption, dosage, and utilization of drug combinations in topical drug delivery[1–3]. Several nanoparticle systems made of polymers, lipids, metal, and dendrimers have been quite successfully utilized as topical delivery vehicles [4–6]. However, one of the key challenges facing transdermal penetrability of nanoparticle is the innate barrier function of skin, which is significantly determined by the overall skin thickness [7,8]. Interestingly, skin thickness is not uniform across the body and tends to vary at different anatomical sites to suit function and mobility. Particularly, skin overlying the palm and feet are much thicker, devoid of hair and differ in tissue morphology, compared to skin from other areas. Future topical strategies for treating conditions that exclusively affect thick skin, such as palmar-plantar erythrodysesthesia and palmo-plantar keratoderma, would need to address the intrinsic morphological differences between thick and thin skin[9,10].

The stratum corneum (SC) layer of the epidermis contributes significantly to skin thickness and is generally considered as an important barrier that needs to be overcome for topical nanoparticle delivery. The SC is primarily made of keratin-filled dead corneocytes embedded in a lipid rich matrix that makes it challenging for transdermal penetration of both hydrophobic and hydrophilic compounds[11,12]. Just below the SC, thick skin contains an additional clear keratohyalin layer called the stratum lucidum, which has also been attributed to reduced penetrability of thick skin.

Over the years, an important strategy to enhance transdermal nanoparticle penetration has been to circumvent the SC via physical disruption of the SC[4,13,14]. Application of an adhesive tape to strip the superficial epidermal layer[15,16], microneedle applied abrasions[17,18], electroporation[19], ultrasound application[20], chemical disruption[21,22], and degradable drug loaded microneedles[23] are few such strategies. However, almost all studies that investigate strategies to bypass the SC, have been in the thin skin. To the best of our knowledge, we are unaware of any study that has examined the role of mechano-physical disruption of the superficial layers of thick skin epidermis to promote nanoparticle penetration after topical delivery. In a previous report, intact thick skin behaved as a depot to topically applied gold nanoparticles, and nanoparticles gradually diffused from the epidermal layers even after the topical exposure had ceased[24]. But it is still unclear how nanoparticles would penetrate thick skin after mechano-physical disruption to the epidermal layers. In this study, we characterize and compare two important mechano-physical strategies of epidermal disruption (tape-stripping and microneedle abrasion) in thick and thin skin. We further asked how such physical disruption of the epidermal surface in thick and thin skin would affect the penetration of topically applied polystyrene nanoparticles.

## 2. Materials and methods

### 2.1. Materials

Fluoresbrite® YO carboxylate microspheres 100nm (Polystyrene nanoparticles) were purchased from Polysciences, USA. L929 mouse fibroblast cell lines were procured from National Centre for Cell Science, Pune. MTT (3-(4,5-dimethylthiazol-2-yl)-2,5-diphenyltetrazolium) reagent and paraformaldehyde were purchased from Sigma-Aldrich, USA. Dulbecco’s Modified Eagle’s Medium (DMEM) was purchased from Lonza Chemicals, USA. Fetal bovine serum (FBS) was purchased from Life technologies, USA. Penicillin and streptomycin mixture was purchased from Invitrogen, USA. 2-Propanol, hydrochloric acid 35% (HCl), sodium chloride, Di-sodium hydrogen phosphate, potassium chloride, potassium dihydrogen phosphate and sodium hydroxide were purchased from Merck Chemicals, USA. Methylene Blue was purchased from Loba Chemie, India. Mili-Q water was used in all experiments.

### 2.2. Animal care

Skin samples utilized in this study were obtained from Sprague-Dawley rats (male or female) weighing 250-325 g that were housed in pairs and allowed standard rat diet and water ad libitum. Cages were maintained on 10 h/14 h light/dark cycle. All animals used in this study were covered under protocols approved by the Institutional Animal Ethics Committee, Amrita Institute of Medical Sciences and Research Center, Kochi, India, in accordance with guidelines set forth by CPCSEA, government of India. Shaved skin from the anterior abdominal wall was used as a model for thin skin, while skin from the central plantar surface area of the hind paw was used as a model for thick skin.

### 2.3. Histological analysis of skin

Skin samples were obtained from rats that were euthanized by carbon dioxide inhalation. Skin segments measuring approximately 1 cm x 1 cm were gently removed from the plantar surface of the hind paw and anterior abdominal wall. Skin segments were fixed in 4% paraformaldehyde after washing with PBS and carefully removing any underlying fat tissue and overlying hair. Samples were subjected to serial sectioning (5 µm thickness) and standard hematoxylin and eosin staining protocols. Sections were cover slipped and imaged under bright field microscopy using DM500 microscope attached to an ICC50 HD camera (Leica, Germany). Epidermal thickness was measured and quantified in three randomly chosen fields from 8 selected images per experimental group using ImageJ software (Ref).

### 2.4. Tape stripping and assessment of tape peeling strength

Superficial layers of skin were removed from thick and thin skin from euthanized animals using the tape stripping method. Briefly, commercially available adhesive tape strips (3.9 cm^2^ surface area) were gently pressed against the skin surface and quickly peeled off at a 45LJ angle. The process was repeated 20 times with a fresh adhesive tape strip each time. Tape stripped skin were further utilized for transdermal studies and histological assessment of skin thickness.

The force required to peel off the adhesive tape from the skin surface was assessed using a universal tensile strength machine (Shimadzu, Japan), at a peeling speed of 100 mm/min. The average force required to peel was analyzed using the Trapezium X software, with each test performed 20 times on each skin sample.

### 2.5. Microneedle application

Thick and thin skin areas were cleaned, and a microneedle-based derma roller (Kostech Derma roller) was applied with back-and-forth movements (10 cycles), along with a gentle downward force. The length of the microneedle within the head of the derma roller was 500 µm. The average number of micro punctures per unit area of skin was quantified from images obtained after running the derma-roller over intact skin.

### 2.6. Intradermal injection of PS nanoparticles

Excised skin from hind paw and abdomen were injected with 25 µL of rhodamine encapsulated PS nanoparticles (1% w/v) suspended in PBS using an insulin needle and syringe. Care was taken to inject the nanoparticle solution within the epidermal-dermal area. A successful intradermal injection resulted in a distinct bleb on the skin surface.

### 2.7. Size characterization of polystyrene nanoparticles

Hydrodynamic particle size distribution and zeta potential of polystyrene nanoparticles were determined using zetasizer Nano ZS (Malvern, UK). Further confirmation of the size of the nanoparticles was determined using Scanning Electron Microscopy (Jeol JSM 649 Analytical Scanning Electron Microscopy, Japan).

### 2.8. Ex-vivo skin permeation assay

Freshly harvested thick and thin skin segments from euthanized rats were cleaned and used for the permeation assay. Skin segments were mounted between the donor and the receptor chamber of a glass diffusion device maintained at 37°C. The chamber was checked for absence of leaks before commencement of experiments. The donor chamber was filled with 2 mL of rhodamine encapsulated PS nanoparticles, while the receptor chamber was filled with distilled water. Absolute care was taken to remove any bubbles that were formed underneath the skin samples. Periodic samples were obtained from the receptor chamber via the sampling port, without disturbing the layout of the skin segment between the chambers. Translocation of nanoparticles across the skin segment was determined by measuring the fluorescence intensity of the collected receptor solution using a microplate UV/visible spectrophotometric plate reader (Bioteck Synergy HT). Each sample was measured in triplicate from at least 3 skin segments.

### 2.9. Statistical analysis

All the data shown are in mean ± standard deviation and statistical comparison between experimental groups were performed using unpaired Student’s t test with multiple comparison where necessary.

## 3. Results

### 3.1. Epidermal thickness of rat hind paw and abdominal skin

The overall thickness of the skin varies according to anatomical site. To specifically assess the thickness of stratum corneum and epidermis in thick and thin skin, we harvested skin samples from rat hind paw and anterior abdominal wall respectively. Skin segment from in-between the foot pads was used for histological assessment of thick skin. Hematoxylin and eosin-stained skin sections demonstrated distinct epidermal and dermal demarcation in both thick and thin skin. The overall thickness of the epidermis in hind paw skin was 115.7 ± 16.4 µm, as compared to 29.2 ± 5.1 µm in the abdominal wall skin (Figure 1), a difference of about 75%. Similarly, the stratum corneum of hind paw skin was 67.44 ± 17.3 µm, while that in the abdominal skin was 9.9 ± 0.6 µm. Interestingly, the stratum corneum formed approximately 58% of the total epidermis in the thick skin, while it comprised of only 33% in the thin skin.

**Figure 1.**
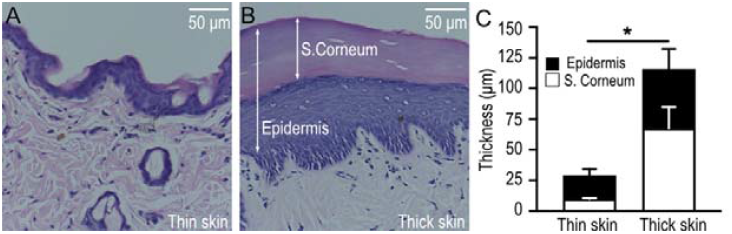
Thickness of epidermis in thin and thick skin of rat. Representative hematoxylin and eosin stained thin (A) and thick (B) skin sections harvested from rat abdomen and hind paw respectively. Bar graphs depicting epidermal and stratum corneum thickness in thin and thick skin. Data shown are mean ± S.D, obtained from three random skin sections. Five non-overlapping regions from each skin section was analyzed and quantified. Scale bar: 50 µm

**Figure 2.**
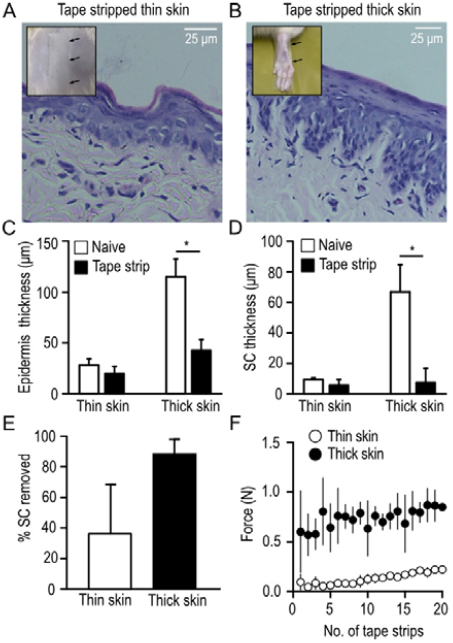
Effect of repetitive tape stripping on epidermal skin thickness. Representative hematoxylin and eosin stained thin (A) and thick (B) skin after subjecting to consecutive tape stripping. A fresh adhesive tape was used for each strip and the process was repeated 20 times. Inset images show the location of tape placement (arrows) on the anterior abdominal wall (A) and hind paw (B). Thickness of tape stripped epidermis (C), stratum corneum (D) and the fraction of stratum corneum removed (E) was quantified and displayed as bar graphs. All data shown are mean ± S.D, obtained from three random skin sections. Scatter plot depicting the force required to ‘strip’ a fresh adhesive tape applied on thin and thick skin, with each data point representing average of 3 independent tape strips (F).

### 3.2. Effect of tape stripping on epidermal thickness in thick and thin skin

To evaluate the impact of repetitive tape stripping on epidermal skin, both hind paw and abdominal wall skin segments were subjected to 20 repetitions of tape stripping. A histological assessment was then performed to measure the epidermal and stratum corneum thicknesses. The results showed that tape-stripping reduced the overall thickness of hind paw epidermal skin from 115.7 ± 16.4 µm to 43.5 ± 9.7 µm, a reduction of 63%. In contrast, tape-stripping reduced the overall thickness of abdominal skin from 29.1 ± 5.1 µm to 21 ± 5.9 µm, a reduction of only 28%. Large reduction in thickness was particularly observed in the stratum corneum. Tape-stripping reduced stratum corneum thickness in hind paw skin from 67.4 ± 17.3 µm to 8.2 ± 8.5 µm, a reduction of almost 87%. However, in the abdominal skin, tape-stripping caused a smaller decrease in stratum corneum thickness, from 9.9 ± 0.6 µm to 6.2 ± 3.2 µm, a reduction of 38%. Histologically, apart from the reduction in stratum corneum thickness, the rest of the epidermal architecture demonstrated normal morphology. To further evaluate the reason behind differential reduction in stratum corneum thickness between thick and thin skin, we measured the force required to peel the tape off the skin using a tensile strength machine. Interestingly, the peeling force for thick skin was 4-5 times higher than that required for thin skin. The required peeling force remained constant throughout the 20 repetitions for both thick and thin skin.

### 3.3. Characterization of microneedle-roller induced puncture wounds on skin

Next, to evaluate the nature of skin aberration induced by a microneedle-roller, we utilized a commercially available microneedle-roller with a needle length of 500 µm as shown in Figure 3A. The roller was applied on a specific area of the skin and the resulting microscopic skin perforations in terms of perforation density and inter-perforation distance were quantified. Results showed that the skin perforation density was 0.15 ± 0.03/mm^2^ and the inter-perforation distance was 1.9 ± 0.09 mm. Closer observation of the micro roller induced skin perforation showed that each perforation had a smooth entry and exit path and demonstrated a uniform distribution pattern.

**Figure 3.**
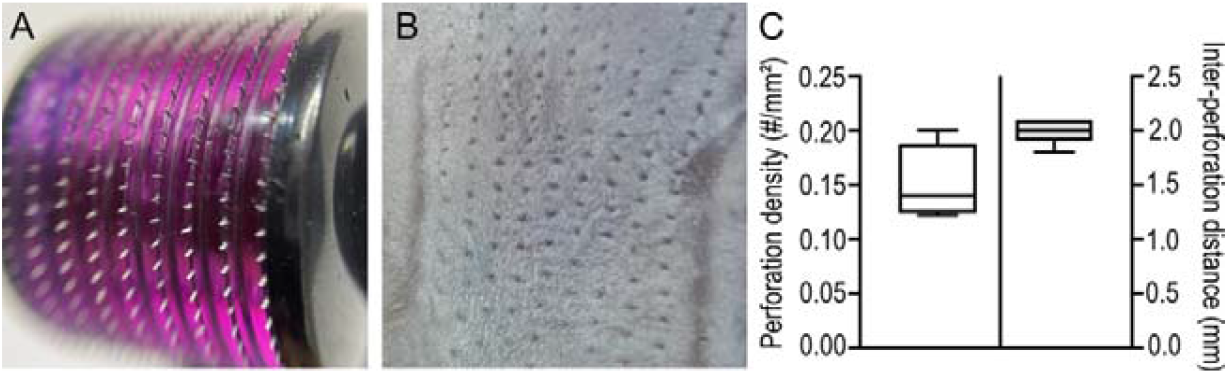
Characterization of microneedle-roller induced perforations. Representative image showing the microneedle roller head used in the study (A), with each needle measuring about 500 μm in length. Perforation density and inter-perforation distance was quantified by applying the microneedle roller intact rat skin (B), and the values were quantified and displayed as box-whisker plot (C). Scale bar: 5 mm.

### 3.4. Intradermal injection of rhodamine encapsulated polymeric nanoparticles in thick and thin skin

To determine the overall penetrability of nanoparticles via the subepidermal layers of thick and thin skin, we deposited nanoparticles in the intradermal space, just underneath the epidermis (Figure 4A). Commercially sourced rhodamine encapsulated polymeric nanoparticles with hydrodynamic size of 122.5 ± 31.35 nm and surface diameter of 127.47 ± 10.64 nm were carefully injected under the epidermis of thick and thin skin (Figure 4D, E). Injected nanoparticles were visible as pinkish stain underneath the skin (Figure B, C).

**Figure 4.**
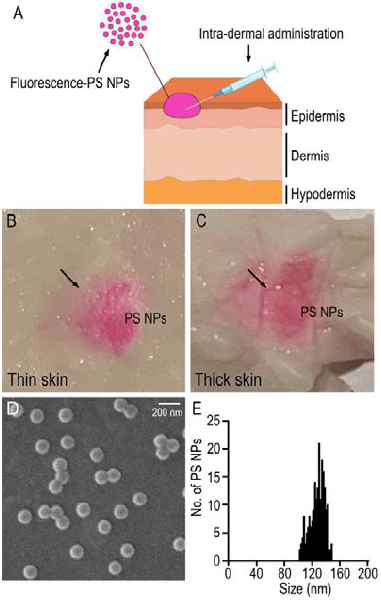
Intradermal injection of rhodamine encapsulated polymeric nanoparticles in thick and thin skin. Illustration showing the injection of polystyrene nanoparticles into the sub-epidermal region of the skin (A) as perfromed in this study. Representative image of thin (B) and thick (C) skin adminstered with intradermal polystyrene nanoparticles, with arrows indicating the bleb formed after the intradermal injection. Representative SEM images of rhodamine encapsulated polystyrene nanoparticles as shown in (D) were quantified and the overall distribution of particle size (diameter) displayed as histogram in (E).

### 3.5. Tape stripping enhances nanoparticle penetration in thick skin

Diffusion of polystyrene nanoparticles across skin subjected to mechano-physical disruption of the epidermal surface was tested in a modified Franz diffusion chamber comprising of a donor and receptor chamber with skin samples placed in between. Skin sample with intradermal inoculation of nanoparticles was included to assess subepidermal diffusion across the skin. Diffusate from the receptor chamber was measured for fluorescence intensity at varying time points. We observed a gradual increase in nanoparticle accumulation in the receptor chamber in all groups tested. Tape stripped hind paw skin showed clear increase in nanoparticle diffusion after 2 to 3 hours compared to non-stripped naïve skin (P<0.05). Curiously, this difference in diffusion rates between tape-stripped and non-stripped skin was absent in skin derived from the abdominal wall (Figure 5A, B). Furthermore, the observed difference in nanoparticle diffusion between tape-stripped and naive hind paw skin disappeared after 24 hours. It was also interesting to note that skin subjected to microneedle punctures showed comparable diffusion rates to that of naïve skin in both thick and thin skin at all time points. The skin segments that were intradermally injected with nanoparticles showed similar nanoparticle diffusion pattern irrespective of the skin type. Results from this experiment demonstrates that tape-stripping was far more effective in promoting nanoparticle penetration in the thick skin as compared to thin skin.

**Figure 5.**
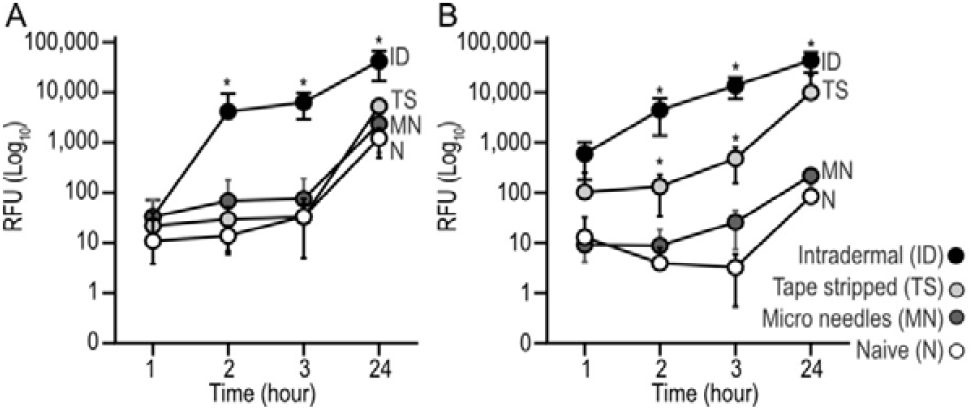
Tape stripping enhances nanoparticle penetration in thick skin. Scatter plot showing fluorescence intensity from the diffusate collected from the diffusion chamber housing either thin or thick (B) skin. Skin samples were either subjected to intradermal injection, tape stripping or microneedle treatment, while naïve skin was used as controls. Each data point represents fluorescent intensity obtained from diffused nanoparticles across three individual skin samples (n=3) per group, at the indicated time points. Data shown as mean ± SD, performed in triplicates, *p < 0.05, One-way ANOVA with multiple comparison test.

## 4. Discussion

The purpose of this study was to investigate the effect of mechanical disruption of the thick skin epidermis on the transdermal penetration of topically applied nanoparticles. The motivation for this work is the current lack of knowledge about nanoparticle transdermal penetration across thick skin types and how disrupting the outer epidermis affects nanoparticle movement. Intriguingly, not many studies have addressed transdermal diffusion of nanoparticles across the thick skin, however, experiments in non-thick skin have revealed that transdermal diffusion of nanoparticles across skin thickness is dependent on both nanoparticle characteristics[21,25– 27], and skin morphology[28,29].

Histologically, skin is composed of the outer epidermis, and the inner dermis. The epidermis is the most important determinant of skin thickness and varies in different locations of the body depending on function. In areas that experience high levels of friction, such as the palms and soles of feet, the epidermis is considerably thicker and is hence referred as the “thick skin.” Stratum corneum, which forms the outermost layer of epidermis is considerably thicker in thick skin and poses a formidable barrier for transdermal penetration of nanoparticles[30,31]. However, in addition to a thicker SC, the thick skin epidermis has an extra layer of keratohyalin called the stratum lucidum (SL) [32,33]. Thin skin is distinguished by the absence of SL. Curiously, the contribution of SL to nanoparticle penetration is not completely clear, although reduced nanoparticle penetrability has been proposed[33,34]. The remaining three layers of the epidermis, viz, stratum spinosum, stratum granulosum and stratum basale are common for thick and thin skin but vary slightly in their overall thickness between skin types. In our study, we confirmed that the rat SC derived from thick skin is approximately 5-6-fold thicker than thin skin. This observation is quite similar to what is seen in humans where thick skin is approximately 3-4 fold thicker than thin skin[35].

Previously, studies investigating the topical application of gold nanoparticles on thick skin found that particles accumulated underneath the SC and within the lower layers of the epidermis[24]. The thick epidermis acted as a depot, and the gold nanoparticles were reported to gradually diffuse across the dermis over time. To further enhance nanoparticle penetration, various forms of mechano-physical disruption of the SC and epidermis have been proposed, including tape stripping and mechanical skin abrasion. While the effectiveness of these techniques in enhancing the penetration of topically applied drug formulations is well-documented[35,36] their effectiveness in aiding nanoparticle penetration across skin is relatively recent[37,38]. Moreover, the contribution of SC in restricting nanoparticle entry and the actual role of mechano-physical disruption in facilitating nanoparticle penetration remains unclear. The limited consensus is primarily because there have been several studies that have yielded starkly contrasting results. For instance, some studies have found that topical application of quantum dots and silica nanoparticles on tape-stripped skin had limited effect on epidermal penetration[39,40], while in another study, QD application across human skin showed enhanced SC penetration after tape-stripping[41]. The reason for such contrasting observations could likely be due to varying experimental conditions, nanoparticle characteristics, or skin types. It is worth noting that almost all studies pertaining to particle-associated skin penetration have been conducted in non-thick skin samples obtained from humans[42,43], rats[44], and pigs[45,46].

In this study, we used the tape stripping and micro perforation methods to disrupt the SC in the thick skin of rats. It was interesting to note that tape stripping removed about 80% of the SC in thick skin as compared to only 35% in thin skin. Curiously, the SC layer that was retained after repetitive tape peeling was surprisingly of similar thickness in both thick and thin skin. The most plausible explanation for this observation, considering that a larger force was required to perform each tape peel in the thick skin, is that a large proportion of SC was removed with each peel. Once 80% of the SC was removed, the remanent SC layers may be too adherent to the underlying epidermal layers for any further removal. A similar phenomenon could explain the remnant SC in the thin skin as well. Furthermore, the reason behind epidermal disruption during tape stripping is the result of interplay between the adhesive force of the tape and the cohesive force among the corneocytes in the SC. The number of corneocytes removed by the adhesive tape reduces with every successive strip, because the corneocytes located in the lower layer of the SC exhibit stronger intercellular connections due to modified desmosomes [16,47]. Our findings align with previous studies and we found that despite multiple tape strips, approximately one third of the SC layer remains intact.

To test skin penetration, we used polystyrene particles due to their homogenous size distribution (approximately 110 nm diameter), non-degradability, and high efficiency in retaining the encapsulated fluorophore. To control for possible resistance to transdermal diffusion by sub-SC layers of the skin, we injected nanoparticles intradermally into skin segments. Although we took utmost care to deposit the nanoparticles just underneath the SC, it is possible that some nanoparticles were injected into deeper layers of the epidermis. None-the-less, our observations still help in understanding the rapid rate of transdermal diffusion of nanoparticles across the dermis.

Tape stripped thick skin showed modest increase in transdermal penetration of topical nanoparticles after 2-3 hours compared to thin skin. However, penetration was comparable between skin types after 24 hours. The difference in nanoparticle penetration at 2 and 3 hours may be attributed to the large percentage of SC that was removed by the adhesive tape, suggesting that tape stripping has a much larger effect on nanoparticle penetration in thick skin, as compared to thin skin.

Unlike the tape stripping technique, application of micro perforations in both thick and thin skin produced only a modest enhancement in nanoparticle penetration, with no statistical support. Our observation is not in alignment with previous studies that demonstrate enhanced diffusion of drugs and liposomes after pretreatment of skin with derma roller[19,48,49]. Interestingly, it has been found that penetration of formulations is dependent on needle size, with shorter needle length (150 microns) promoting better penetration[50]. In our study, the needle length was of medium length (500 microns), which might have contributed to the limited nanoparticle penetration.

## 5. Conclusion

In conclusion, we observed that tape stripping facilitated the penetration of polymeric nanoparticles across thick skin, while micro needle application was not effective. We observed that both strategies were ineffective in promoting polymeric nanoparticle penetration in the non-thick skin. Findings of this study could facilitate future drug delivery strategies to treat conditions affecting the thick skin.

## Acknowledgements

The authors would like to thank Dr. A K K Unni, and Dr. Thennavan, Reshmi P with assistance in animal experiments, Department of Polymer Science and Rubber Technology (PSRT) Sophisticated test and Instrumentation Centre (STIC), Cochin University of Science and technology (CUSAT), Kochi for peel test analysis, and Dr. Pallavi Madhusudanan for valuable discussions, experimental assistance and feedback.

## Data availability

All data depicted in this article, including raw data from replicate experiments, analysis and statistical testing have been uploaded into a freely accessible repository. This information can be accessed at the following link: https://doi.org/10.6084/m9.figshare.c.6615751.v1

## Author contribution

**Andrea Antony**: Investigation, methodology, data curation, writing-original draft; **Gayathri Raju**: investigation, data curation; **Ahina Job** and **Meet Joshi**: investigation; **Sahadev Shankarappa:** conceptualization, supervision, project administration, funding acquisition, writing-review & editing, and formal analysis. The work reported in the paper has been performed by the authors, unless clearly specified in the text.

---

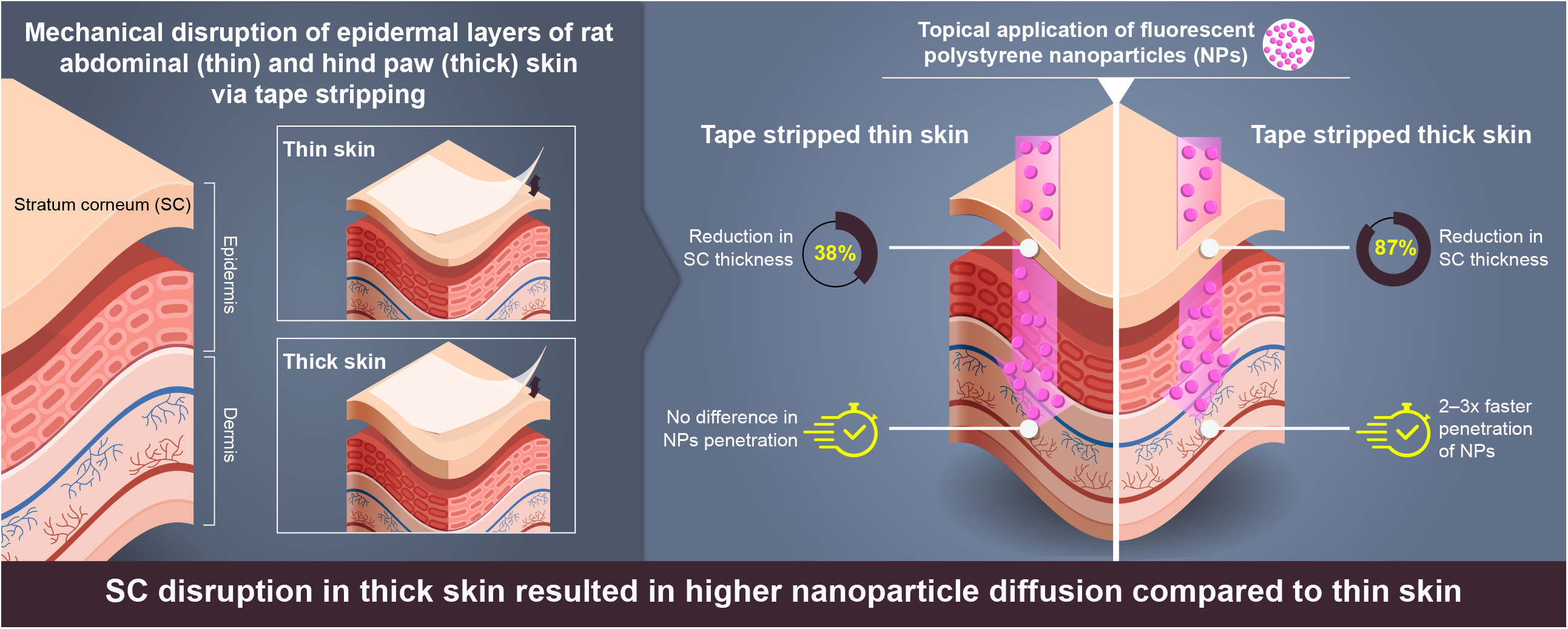

